# Incomplete Reverse Remodeling of the Tricuspid Valve Leaflets Following Relief of Pressure Overload

**DOI:** 10.64898/2026.08.04.742903

**Authors:** Boguslaw Gaweda, Austin Goodyke, Jeremy Prokop, Sanjana Arora, Magda Piekarska, Tomasz A. Timek

## Abstract

**Objective(s):** Tricuspid valve (TV) remodeling and functional tricuspid regurgitation (FTR) progression during right ventricular (RV) pressure overload and reverse remodeling after resolution of RV afterload is poorly understood. We set out to investigate tricuspid leaflet tissue response to induction and subsequent alleviation of pressure overload in a large animal model of RV failure with FTR.

**Methods:** Fifteen healthy adult male Dorset sheep (72±4 kg) underwent pulmonary artery banding (PAB) to induce RV failure and FTR. After 8 weeks, 7 sheep (**PAB**, n=7) were terminated, and remaining 8 had the PAB removed (**rPAB**, n=8) and were followed for another 8 weeks before termination. Both groups underwent epicardial echocardiography and hemodynamic assessment during banding surgery and at terminal operation. Ten healthy sheep served as a control group (**CTL**, n=10) and underwent terminal procedure only. In all animals, TV leaflets and right ventricular (RV) tissue were harvested at terminal procedure and analyzed histologically and transcriptionally.

**Results:** TV leaflets in PAB animals showed increased cross-sectional area and ECM alterations, some of which persisted after resolution of RV pressure overload. rPAB valves exhibited distinct ECM composition, with notably altered mucin and fibrin content, suggesting a shift toward matrix stabilization, dissimilar to control and PAB. RNA sequencing uncovered a unique molecular state in rPAB valves, with persistent changes in PRG4, PDE3A, CXCL8, and HLA transcripts. RV tissue also demonstrated a separate remodeling trajectory, with sustained expression of stress-related genes including PDE3A, NAV2, ANFB, and ACTS. These findings indicate that both valve and ventricular tissues retain a persistent remodeled phenotype post-unloading.

**Conclusions:** TV leaflets actively remodel in response to hemodynamic stress and do not fully revert to a normal state after relief of pressure overload. This persistent altered phenotype may represent a biological contribution of the TV leaflets to recurrent TR with implications for long-term outcomes following treatment of FTR.

*Clinical Perspective:* 

*What is new?:*

- Relief of right ventricular pressure overload, in a large animal model, resulted in substantial reverse remodeling of the right heart and reduction of tricuspid regurgitation severity, but tricuspid valve leaflets did not return to a normal state.
- Reverse remodeled leaflets remained enlarged despite normalization of hemodynamics with an altered extracellular matrix.
- Cellular proliferation and immune cell infiltration observed during pressure overload resolved after unloading, yet transcriptomic analysis identified a distinct molecular phenotype that differed from both healthy and diseased valves.
- Tricuspid valve leaflets are active biological participants in the remodeling process and exhibit persistent adaptation or maladaptation after resolution of the initiating hemodynamic stress.

*What Are the Clinical Implications?:*

- Secondary tricuspid regurgitation should be considered a disease involving both right heart geometry and leaflet biology.
- Resolution of the underlying cause of tricuspid regurgitation may not restore leaflet structure and molecular homeostasis.
- Persistent leaflet remodeling may contribute to residual or recurrent tricuspid regurgitation despite successful treatment of pulmonary hypertension or other inciting conditions.
- Therapies directed at leaflet remodeling may ultimately complement surgical and transcatheter strategies currently focused on annular and ventricular geometry.

## Introduction

Functional tricuspid regurgitation (FTR) is most often secondary to left-sided heart disease or atrial fibrillation where pressure and/or volume overload drive right ventricular (RV) remodeling and/or atrial and tricuspid valve (TV) annular dilation leading to geometrical alterations to the TV apparatus and subsequent incompetence.^1^ Historically, functional TR is indirectly addressed through etiology based approach to surgically correct only the left side, which has been burdened with significant residual and progressive FTR resulting in increased morbidity and mortality.^2,3^ However, when severe FTR is present, even surgical correction with the contemporary gold standard of prosthetic ring annuloplasty produces suboptimal results.^4,5^ Although significant progress has been made in our understanding of the complex geometric and hemodynamic changes leading to FTR, minimal consideration has been given to leaflet tissue in the pathogenesis and progression of FTR. Despite increasing evidence from pre-clinical and clinical studies suggesting leaflet tissue remodeling^6,7^ or reverse remodeling,^8^ the TV leaflets have been thought to be only passive responders to ventricular mechanical and hemodynamic forces, not contributing in the development, progression and regression of TV incompetence. Clearly, FTR is a complex entity with different phenotypes already distinguished by alteration in atrial and ventricular geometry,^9^ but the exact role of the leaflets remains obscure and unaccounted for in treatment.

Using a clinically relevant ovine model of pulmonary artery banding (PAB) induced FTR, we previously demonstrated that TV leaflets undergo significant cellular, extracellular matrix, and transcriptional remodeling in response to chronic pressure overload.^10–12^

However, whether these alterations regress following relief of the hemodynamic burden remains unknown. Understanding this process is clinically important because residual and recurrent TR frequently occur despite successful treatment of the underlying disease process and may reflect persistent valve abnormalities. Therefore, we sought to characterize structural, cellular, and transcriptional changes in tricuspid valve leaflets following reversal of PAB induced pressure overload and FTR. We hypothesized that leaflet reverse remodeling would be incomplete, resulting in a biologically distinct phenotype that differs from both healthy and diseased valves, potentially explaining clinical observations of residual TR, or TR recurrence in the context of resolved primary disease.

## Materials and Methods

All animals were provided humane care in accordance with the Principles of Laboratory Animal Care established by the National Society for Medical Research. The study protocol received approval from the Michigan State University Institutional Animal Care and Use Committee, protocol number PROTO202100330 approved on 01/31/2022. Animals were housed and maintained at the large animal facility of Michigan State University.

### Surgical outline

The utilized surgical protocol has been previously detailed by our group^13^ and will be summarized here briefly. The surgical procedure and experimental groups are schematically outlined in Figure 1. Animals were divided into three groups for the study, those sacrificed without any intervention that served as control (CTL, N=10), those monitored for 8 weeks following PAB (PAB, N=7) and those having the PAB removed after 8 weeks and monitored for an additional 8 weeks (rPAB, N=8).

**Figure 1.**
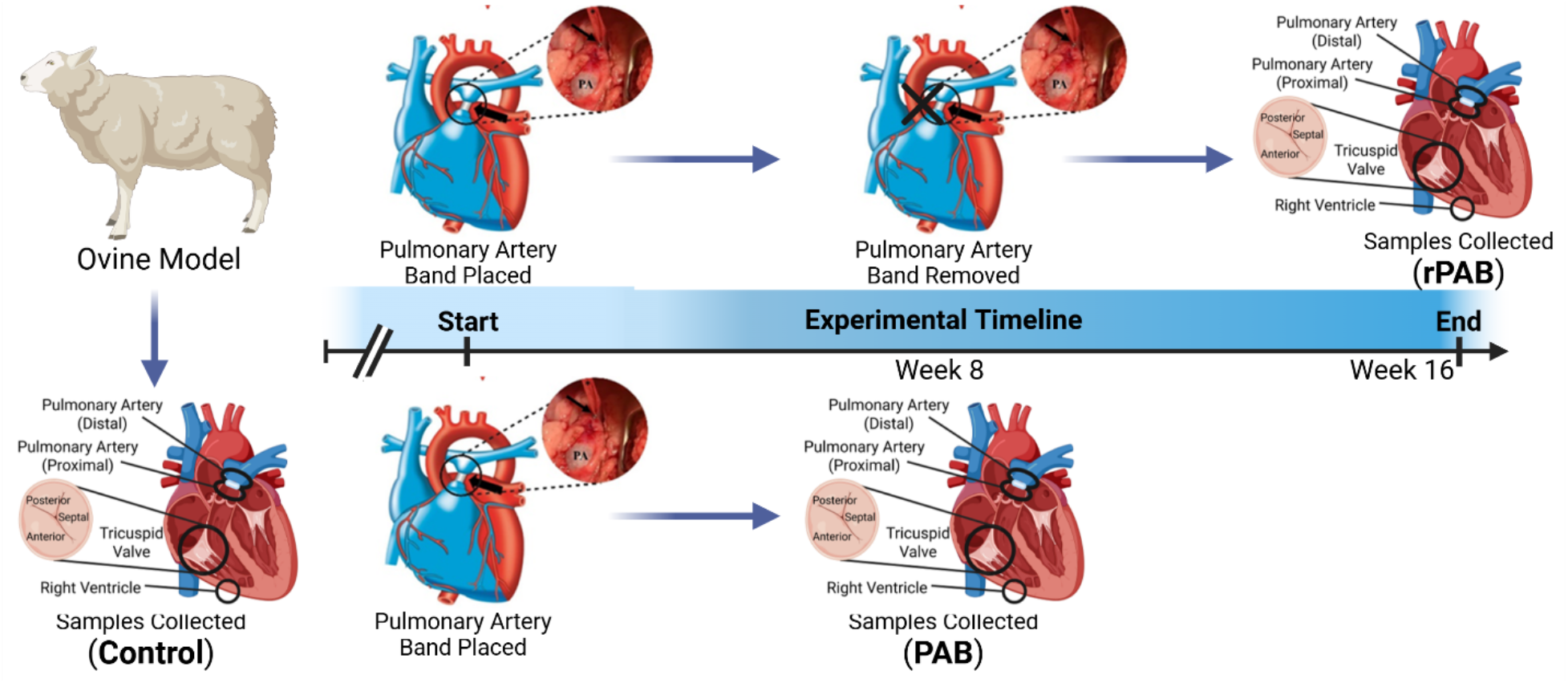
Experimental outline diagraming the pulmonary artery banding procedure, the acquisition of samples (distal and proximal pulmonary arteries; anterior, posterior, and septal leaflets of the tricuspid valve; and the right ventricle), and the timeline for experimental procedures.

### Pulmonary Artery Banding

For the pulmonary artery banding procedure, fifteen healthy adult male Dorset sheep (72±4 kg) underwent placement of an external right jugular intravenous catheter under local anesthesia, utilizing 1% lidocaine for subcutaneous injection. Animals were anesthetized using propofol at a dosage of 2-5 mg/kg intravenously, followed by intubation and mechanical ventilation. General anesthesia was sustained using inhalational isoflurane (1-2.5%) with a fentanyl infusion (5-20 mcg/kg/min) for additional maintenance. A sterile limited left thoracotomy was performed via the 4th intercostal space, followed by epicardial echocardiography to evaluate biventricular function and the competence of the tricuspid and mitral valves. Arterial catheters were placed into left internal mammary artery and pulmonary artery (PA) for hemodynamic monitoring. The main pulmonary artery was subsequently encircled with umbilical tape proximal to its bifurcation. Systemic and pulmonary pressures were monitored while the umbilical tape was progressively tightened using clip approximations until the threshold of hemodynamic stability. Animals were recovered after closure of thoracotomy.

### Pulmonary Artery Debanding

Eight weeks following the PAB procedure, eight animals designated to the rPAB group returned to the operating room for removal of the band. For hemodynamic monitoring, the right radial artery in the forelimb was cannulated. The animals underwent a left re-thoracotomy, the PA band was identified, and a pressure catheter was placed into the PA to record pressures proximally and distally to the PA band. Subsequently, the band was dissected from the pulmonary artery and removed. A custom-made pressure balloon catheter 24x30mm (Nordson Medical Inc Salem, NH) was then introduced into proximal PA and positioned across the stenotic segment. Proper alignment was confirmed manually, and the balloon was inflated several times (5 to 8 atm) until there was no visible external constriction. The success of PA expansion was confirmed by pressure measurements (no gradient along PA) and perivascular ultrasonography (no PA intraluminal stenosis).

### Terminal Procedure

All study animals in the three groups (CTL, PAB, rPAB) underwent median sternotomy at the time of the terminal procedure, followed by epicardial echocardiography to evaluate biventricular function and valvular competence. Pulmonary artery pressures were measured directly with a pressure catheter. At the end of the experiment, the animals were euthanized via the administration of sodium pentothal (100 mg/kg IV). The heart was excised, and each tricuspid leaflet was promptly harvested, with tissue allocated for RNA sequencing and histological analysis.

### Acquisition of echocardiographic data

Epicardial echocardiography (2-4 MHz transducer, Vivid S6, GE Healthcare, USA) was conducted in all animals to evaluate biventricular function, size, and valvular competence. Dimensions and functionality of the RV were determined using apical four-chamber and targeted views. RV fractional area change (RV FAC) was defined as the difference between end-diastolic area and end-systolic area, divided by the end-diastolic area. TAPSE (tricuspid annular plane systolic excursion) was assessed via M-mode from an apical four-chamber view. Tricuspid regurgitation (TR) grading involved a thorough assessment of color flow and continuous-wave Doppler, classified by an echo sonographer into categories: none or trace (0), mild (+1), moderate (+2), moderately severe (+3), or severe (+4).

### RNA-seq

Right ventricle samples as well as leaflets of the tricuspid valve were separated following excision and stored in RNALater (Thermo Fisher), incubated at 4°C overnight, and subsequently frozen at -80°C. RNA was extracted using TRIzol (Invitrogen) and cleaned up with the RNeasy MinElute kit (Qiagen). RNA with RIN scores >7 as determined by a bioanalyzer was used to produce stranded total RNA libraries that were prepared with ribosomal RNA depletion for paired-end 150 bp sequencing (average depth: 50M reads) on a NovaSeq 6000 (Illumina) at Azenta Life Sciences (South Plainfield, NJ). FASTQ files were deposited in the NCBI SRA under BioProject PRJNA1182691 and assessed for quality using FastQC. Reads were aligned to a *de novo* transcriptome assembly, previously generated using 39 paired-end datasets with Trinity,^14^ followed by open reading frame (ORF) prediction using TransDecoder and amino acid annotation via Trinotate Annotation files are available at: https://doi.org/10.6084/m9.figshare.27228696.v1. Principal Component Analysis (PCA) was performed in NetworkAnalyst. Significant transcripts were identified based on expression in at least one sample at ≥1 TPM, with a log2 fold change ≥1 and Bonferroni-corrected p-value <0.05. Heatmaps were generated using Morpheus (Broad Institute) (https://software.broadinstitute.org/morpheus).

### Histology

Tricuspid valve leaflets were individually excised and fixed in 10% formalin for 24–72 hours. Samples were then dehydrated in alcohol and paraffin-infiltrated using a Tissue-Tek VIP Processor. Medial leaflet tissue was embedded on end in paraffin blocks and transversely sectioned at 5 µm by the Corewell Health Histology Core (Grand Rapids, MI), or by the University of Michigan Histology Core (Ann Arbor, MI). Sections were stained with hematoxylin and eosin (H&E) and Movat’s pentachrome or Ki67 and CD45 by the Corewell Health and University of Michigan histology cores respectively. Slides were scanned using a Leica Aperio AT2 Scanner and imaged via Aperio eSlideManager using Aperio ImageScope (Leica Biosystems). Image analysis and quantification of size, collagen, GAGs, elastin, cellularity, Ki67, and CD45 were conducted using ImageJ (v2.1.0/1.53c/Java 1.8.0_172).

### Statistical analysis

Statistical analyses were performed using MedCalc (v20.114). Echocardiographic and histological data were analyzed using two-way ANOVA with Bonferroni correction for all pairwise comparisons. The two factors included in the model were surgical treatment group (CTL, PAB, rPAB) and anatomical leaflet location. Data sets that violated assumptions of normality or homogeneity of variance were log-transformed, and if assumptions remained unmet, the non-parametric Kruskal-Wallis test was applied, followed by Conover post hoc tests for pairwise comparisons. A p-value < 0.05 was considered statistically significant. Error bars represent the standard error of the mean (SEM).

## Results

### Echocardiographic and hemodynamic assessments

No significant differences were observed at baseline in control and interventional groups (PAB, rPAB) in echocardiographic and hemodynamic metrics (Table 1), and similarly, there were no differences in pulmonary artery pressures after banding between PAB and rPAB groups at baseline and after 8 weeks (Table 2). Immediately after banding, systolic pulmonary artery pressure (sPAP) increased significantly (tripled) in PAB and rPAB animals. After 8 weeks, sPAP decreased significantly compared to sPAP measured immediately after the initial banding procedure in both PAB and rPAB animals, with no significant difference between the two groups (Table 2). At 8 weeks, after band removal in rPAB animals, sPAP was immediately reduced and did not differ significantly from baseline and persisted unchanged until the 16-week terminal surgery (Table 3). After 8 weeks of PAB, hemodynamic and echocardiographic metrics were assessed for both PAB and rPAB animals. RVFAC decreased from 52±7 to 29±13 mmHg (p<0.001) and from 51±4 to 21±8 mmHg (p<0.001), and TR (0-4+) increased from 0.4±0.5 to 3.7±0.5 (p<0.001), and from 0.5±0.5 to 3.2±1 in the PAB and rPAB group, respectively and there were no significant differences in either echocardiographic metrics or pulmonary artery pressures at 8 weeks between both groups (Supplemental Table 1).

**Table 1.** Baseline hemodynamic and echocardiographic parameters in all groups.

| Baseline parameters | CTL | PAB | rPAB | p-value |
| --- | --- | --- | --- | --- |
| HR (beats/min) | 96 ± 15 | 108 ± 15 | 99 ± 9 | 0.199 |
| Weight (kg) | 68 ± 5 | 74 ± 4 | 71 ± 4 | 0.051 |
| TR (0-4+) | 0.6 ± 0.5 | 0.4 ± 0.5 | 0.5 ± 0.5 | 0.799 |
| TA (cm) | 2.3 ± 0.5 | 2.5 ± 0.3 | 2.7 ± 0.3 | 0.114 |
| RA area (cm) | 10 ± 3 | 12 ± 3 | 13 ± 1 | 0.143 |
| RVFAC (%) | 56 ± 6 | 52 ± 7 | 51 ± 4 | 0.223 |
| LVEF (%) | 57 ± 6 | 61 ± 4 | 62 ± 6 | 0.101 |
| MR (0-4+) | 0.5 ± 0.5 | 0.2 ± 0.4 | 0.1 ± 0.4 | 0.179 |
| sPAP (mmHg) | 17 ± 3 | 17 ± 2 | 18 ± 5 | 0.871 |

Mean ± SD, CTL=controls, PAB=banded animals, rPAB=debanded animals, HR=heart rate, TR=tricuspid regurgitation, TA=tricuspid annulus size, RA=right atrium, RVFAC=right ventricular fractional area change, MR=mitral regurgitation, sPAP=systolic pulmonary artery pressure, p-value by One-way ANOVA with Bonferroni correction.

**Table 2.** Baseline pre- and post-banding, and 8 weeks pulmonary artery pressures and echocardiographic parameters in banded and debanded groups.

| Baseline parameters | PAB | rPAB | p-value |
| --- | --- | --- | --- |
| sPAP pre-band (mmHg) | 17 ± 3 | 18 ± 5 | 0.629 |
| sPAP post-band (mmHg) | 56 ± 6 | 56 ± 7 | 0.988 |
| Parameters at 8 weeks | PAB | rPAB | p-value |
| sPAP (mmHg) | 39 ± 10 | 39 ± 9 | 0.994 |
| HR (beats/min) | 120 ± 35 | 135 ± 19 | 0.286 |
| Weight (kg) | 79 ± 3 | 76 ± 5 | 0.171 |
| TR (0-4+) | 3.7 ± 0.5 | 3.2 ± 1 | 0.178 |
| TA (cm) | 3.9 ± 0.4 | 4 ± 0.6 | 0.530 |
| RA area (cm) | 29 ± 5 | 23 ± 5 | 0.037* |
| RVFAC (%) | 29 ± 13 | 21 ± 8 | 0.212 |
| LVEF (%) | 53 ± 4 | 60 ± 7 | 0.250 |
| MR (0-4+) | 0.5 ± 0.5 | 0.5 ± 0.7 | 1.00 |

**Table 3.** Echocardiographic and hemodynamic parameters in debanded animals (rPAB group) at studied timepoints.

| Parameters | Baseline | 8 weeks | 16 weeks | p-value |
| --- | --- | --- | --- | --- |
| HR (beats/min) | 99 ± 9 | 135 ± 19* | 105 ± 17<sup>†</sup> | <0.001 |
| Weight (kg) | 71 ± 4 | 76 ± 5 | 81 ± 5<sup>†</sup> | <0.001 |
| RVFAC (%) | 51 ± 4 | 21 ± 8* | 51 ± 10<sup>†</sup> | <0.001 |
| LVEF (%) | 62 ± 6 | 60 ± 7 | 58 ± 7 | 0.499 |
| TA size (cm) | 2.7 ± 0.3 | 4 ± 0.6* | 3.4 ± 0.6* | <0.001 |
| RVD1 (cm) | 2.9 ± 0.3 | 4.6 ± 0.6* | 3.1 ± 0.3<sup>†</sup> | <0.001 |
| RVD2 (cm) | 2.6 ± 0.5 | 4.1 ± 0.7* | 2.4 ± 0.5<sup>†</sup> | <0.001 |
| RA area (cm) | 13 ± 1 | 23 ± 5* | 16 ± 4<sup>†</sup> | <0.001 |
| MR (0-4+) | 0.1 ± 0.4 | 0.5 ± 0.7 | 0.3 ± 0.5 | 0.543 |
| TR (0-4+) | 0.5 ± 0.5 | 3.2 ± 1* | 1.1 ± 1.2*<sup>†</sup> | <0.001 |

| Parameters | pre-band | post-band | pre-deband | post-deband | terminal | p-value |
| --- | --- | --- | --- | --- | --- | --- |
| sPAP (mmHg) | 18 ± 5 | 56 ± 7* | 39 ± 9*<sup>‡</sup> | 17 ± 5<sup>†‡</sup> | 18 ± 5<sup>†‡</sup> | <0.001 |

Mean ± SD, HR=heart rate, RVFAC=right ventricular fractional area change, TA=tricuspid annulus, RVD1=right ventricular basal diameter, RVD2=right ventricular mid-cavity diameter, RA=right atrium, MR=mitral regurgitation, TR=tricuspid regurgitation, PAP=pulmonary artery pressure\* p-value <0.05 vs Baseline, <sup>†</sup> p-value <0.05 vs 8 weeks by One-way ANOVA with Bonferroni correction. Mean ± SD, sPAP=systolic pulmonary artery pressure, \* p-value <0.05 vs Baseline pre-band, <sup>†</sup> p-value <0.05 vs 8 weeks pre-deband, <sup>‡</sup> p-value <0.05 vs Baseline post-band by One-way ANOVA with Bonferroni correction.

At the terminal surgery and time of sample collection (Table 4), there were no significant difference in heart rate between all groups, weight for PAB and rPAB animals was significantly higher than in control animals but not different from each other. TR grade for PAB animals (3.7±0.5) was significantly higher than controls (0.6±0.5) and rPAB animals (1.1±1.2) (p<0.001) and there was no significant difference between controls and rPAB. Echocardiographic metrics of RV geometry revealed a significant decrease in RVFAC and an increase in RVD2 for PAB animals in comparison to both controls and rPAB groups with no differences between the rPAB and controls (RVFAC p<0.001, RVD2 p<0.001). RA area and RVD1, however, were significantly increased in PAB and rPAB animals in comparison to controls, with rPAB animals significantly smaller than PAB (RA area p<0.001, RVD1 p<0.001). There were no significant differences in MR or left heart geometry between all respective groups.

**Table 4.** Terminal echocardiographic and hemodynamic parameters in all groups.

| Parameters | CTL | PAB | rPAB | p-value ANOVA |
| --- | --- | --- | --- | --- |
| HR | 96 ± 15 | 120 ± 35 | 105 ± 17 | 0.127 |
| Weight (kg) | 68 ± 6 | 79 ± 3* | 81 ± 5* | <0.001 |
| TR | 0.6 ± 0.5 | 3.7 ± 0.5* | 1.1 ± 1.2<sup>†</sup> | <0.001 |
| TA (cm) | 2.3 ± 0.5 | 3.9 ± 0.4* | 3.4 ± 0.6* | <0.001 |
| RA area (cm) | 10.3 ± 3.2 | 29 ± 4.8* | 16 ± 3.6*<sup>†</sup> | <0.001 |
| RVFAC (%) | 56 ± 6.6 | 29 ± 3.4* | 51 ± 10<sup>†</sup> | <0.001 |
| RVFWd (cm) | 5.9 ± 0.8 | 5.5 ± 0.9 | 6.2 ± 0.7 | 0.311 |
| RVD1 (cm) | 2.5 ± 0.5 | 4.3 ± 0.5* | 3.1 ± 0.3*<sup>†</sup> | <0.001 |
| RVD2 (cm) | 1.8 ± 0.5 | 3.7 ± 1* | 2.4 ± 0.5<sup>†</sup> | <0.001 |
| RVD3 (cm) | 4.9 ± 0.6 | 5.9 ± 0.3* | 5.5 ± 0.6 | 0.006 |
| LVEF | 55 ± 6 | 53 ± 13 | 58 ± 7 | 0.464 |
| MR | 0.5 ± 0.5 | 0.5 ± 0.5 | 0.3 ± 0.5 | 0.668 |
| LVIDd | 4.7 ± 0.5 | 4.4 ± 1 | 4.5 ± 0.4 | 0.824 |
| LVISd | 3.6 ± 0.6 | 3.4 ± 0.8 | 3.4 ± 0.3 | 0.867 |
| TAPSE | 1.4 ± 0.3 | 0.9 ± 0.4* | 1.5 ± 0.4<sup>†</sup> | 0.014 |
| sPAP | 17 ± 3 | 39 ± 10* | 18 ± 5<sup>†</sup> | <0.001 |

Mean ± SD, PAB=banded group, rPAB=debanded group, HR=heart rate, TR=tricuspid regurgitation, TA=tricuspid annulus, RA=right atrium, RVFAC=right ventricular fractional area change, RVFWd= RV free wall diameter, RVD1= basal RV diameter, RVD2= mid RV diameter, RVD3=RV longitudinal diameter, MR=mitral regurgitation, LVIDd=LV internal diastolic diameter, LVISd=LV internal systolic diameter, TAPSE=tricuspid annulus plane systolic excursion, sPAP =systolic pulmonary artery pressure, \* p-value <0.05 vs CTL, <sup>†</sup> p-value <0.05 vs Banded group by One-way ANOVA with Bonferroni correction.

### TV Leaflet Histology

Our group has previously reported alterations in leaflet size due to PAB in an 8-week and 16-week models of ovine FTR.^12,15^ Figure 2A demonstrates the fold change in leaflet area across the anterior (ATL), posterior (PTL), and septal (STL) leaflets at the annulus, belly, and free edge regions with significant increases in the PAB (p<0.001) and rPAB (p-0.005) animals in comparison to control. The annulus, belly, and free edge were defined as approximate regions as annulus roughly splitting the transverse section of the leaflets into thirds. A Movat’s pentachrome stain was applied to the leaflets demonstrating a trend towards decreased collagen in the PAB animals and an increase in the rPAB animals (Figure 2B), though due to heterogenous responses this trend failed to reach significance (p-0.131). Mucin (Figure 2C) demonstrated a significant increase in the PAB animals (p- 0.001) with a significant decrease in the rPAB animals (p<0.001) compared to PAB (p- 0.0512 compared to control) and showed a significantly different response when comparing the STL and ATL (p-0.0177). A significant increase in fibrin (Figure 2D) was observed in the rPAB animals (p-0.0105) was observed, with noticeable fold changes in the belly and free edges of the ATL and PTL. Increases in nuclei counts (Figure 2E) were observed in both the PAB (p<0.001) and rPAB animals (p-001), though there was no significant difference in cellularity as defined by the number of nuclei per area (Figure 2F). Representative pentachrome stained images of control (Figure 2F), PAB (Figure 2G), and rPAB (Figure 2H) ATL demonstrate the observed alterations in size, mucin, and fibrin.

**Figure 2.**
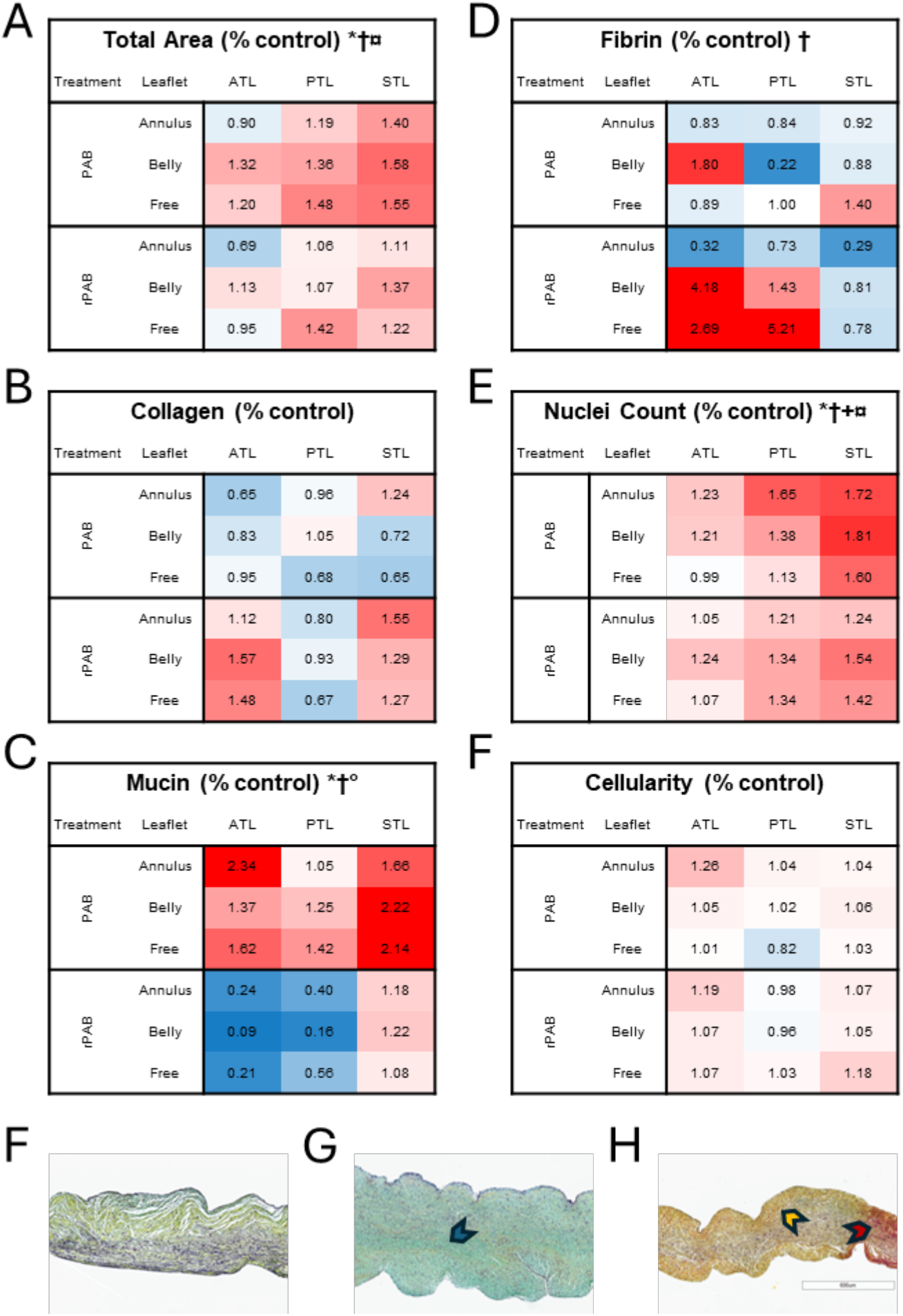
Heat maps showing fold change differences in area (A), collagen (B), mucin (C), fibrin (D), as well as nuclei counts (E) and cellularity (F) measured from cross sectional Movat’s pentachrome stained tricuspid valve leaflets. Anterior, posterior, and septal leaflets are shown with relative fold change differences from control at the annulus, belly and free edge regions of the leaflets. Representative pentachrome stained images of control (F), PAB (G), and rPAB (H) leaflets demonstrate increases in mucin (blue arrow) in the PAB leaflets as well as collagen (yellow arrow) and fibrin (red arrow) in the rPAB leaflets. ATL = anterior tricuspid leaflet, PTL = posterior tricuspid leaflet, STL = septal tricuspid leaflet, * - p<0.05 control vs PAB, † - p<0.05 control vs rPAB, + - p<0.05 PAB vs rPAB, ° - p<0.05 PTL vs STL, ¤ - p<0.05 ATL vs STL.

Leaflets from control, PAB, and rPAB animals were stained for CD45 (Figure 3A) and Ki67 (Figure 3B). Immune cell composition of the leaflet, measured by CD45 (Figure 3C) revealed a significant difference between CTL, PAB, and rPAB groups (p=0.001). *Post hoc* analysis demonstrated a significant increase in CD45 cells in the PAB group in comparison to CTL (p=0.044) and rPAB (p=0.001). Proliferation, measured by ki67 (Figure 3D), demonstrated a similar pattern with a significant difference between groups (p<0.001) as well as a significant difference between leaflets (p=0.017). A significant increase in ki67 was observed in the PAB group in comparison to CTL (p<0.001) and rPAB (p<0.001). The septal leaflet also demonstrated significantly elevated ki67 in comparison to posterior leaflets (p=0.016). Neither CD45 nor ki67 demonstrated any significant differences between CTL and rPAB groups and interaction terms were not significant.

**Figure 3.**
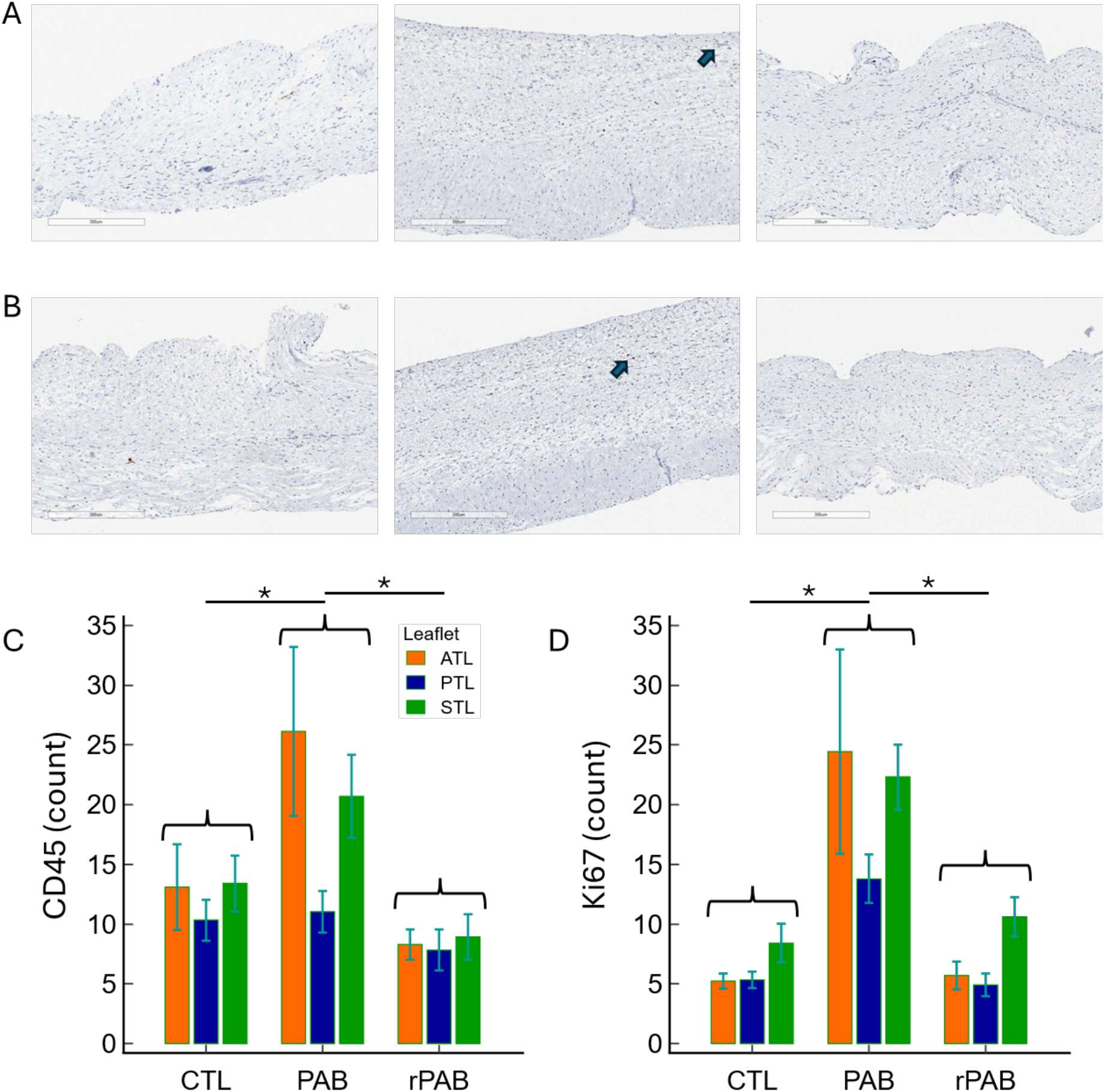
Histologic assessment and quantitative analysis of marker expression across valve leaflets. Representative immunohistochemical staining images of CD45 (A) and Ki67 (B) from anterior leaflets of control (left), PAB (middle), and rPAB (right). Brown chromogenic staining indicates positive immunoreactivity, with arrows highlighting representative positively stained regions. Scale bars = 300 μm. CD45 counts were increased in PAB animals in comparison ot controls and rPAB (C). Ki67 counts increased in PAB in comparison to both control and rPAB (D). Bars represent mean ± SD. ATL = anterior leaflet; PTL = posterior leaflet; STL = septal leaflet; CTL = control; PAB = pulmonary artery banding; rPAB = reverserd pulmonary artery banding; * = p < 0.05.

### Transcriptional Analysis

RNA sequencing was used to assess transcriptional alterations in gene expression between the RV and TV leaflets in control, PAB, and rPAB animals. Pearson minus one clustering of differentially expressed transcripts across the control, PAB, and rPAB conditions for RV and respective leaflets of the TV (Figure 4A) showed clear differential expression patterns between the treatment conditions. Treatment condition and tissue type are shown to be key drivers of the differential expression, with minimal differences between the leaflets. Several transcripts demonstrating these expression patterns are highlighted (Figure 4B), including those associated with endocrine signaling (ANFB), cardiac (ACTS) and leaflet tissue (PRG4) remodeling, cardiac cyclic nucleotide regulation (PDE3A), and immune regulation (HA1A, HA1B).

**Figure 4.**
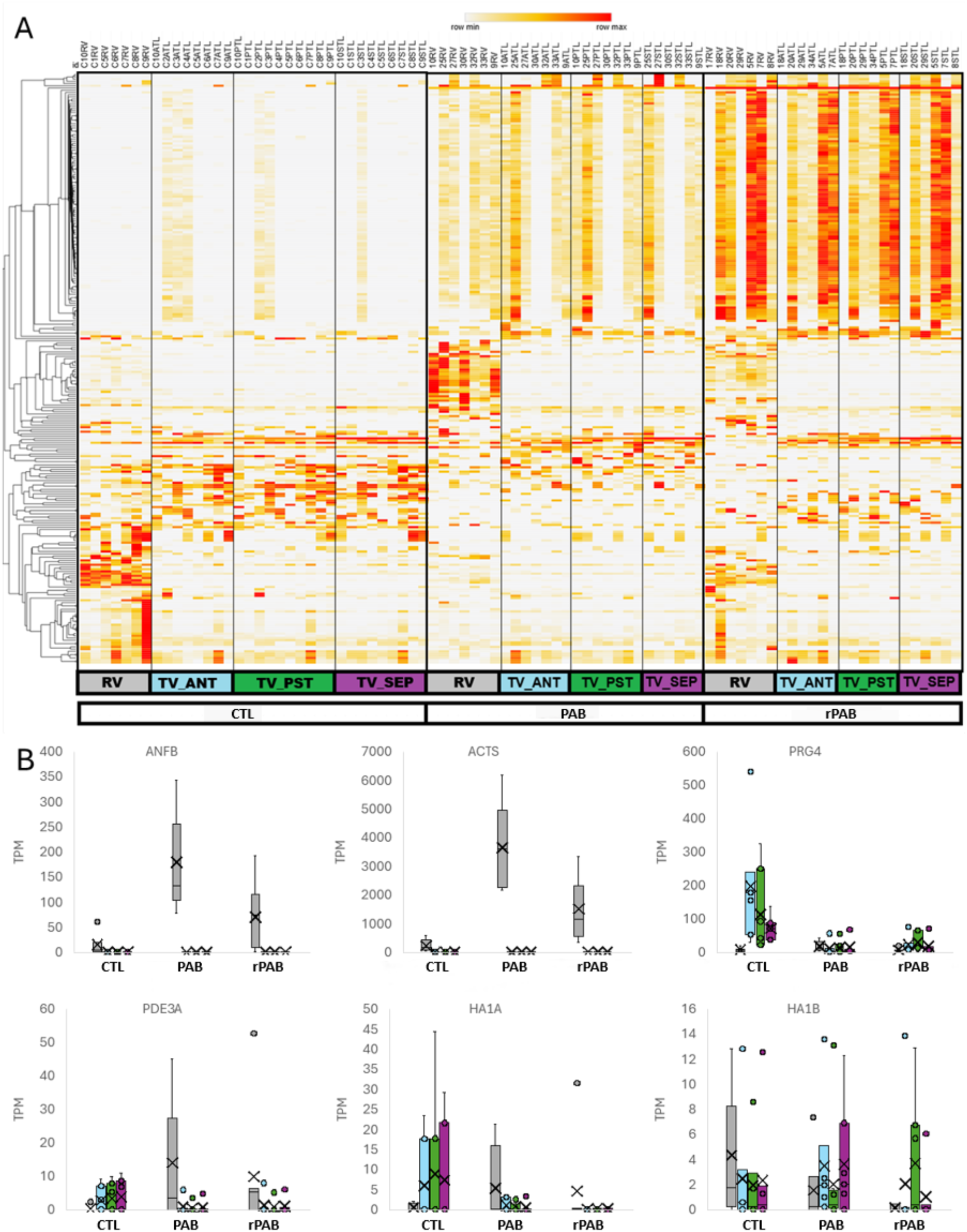
Heat map demonstrating expression differences between control, PAB, and rPAB tissue for right ventricle and the anterior, posterior, and septal tricuspid leaflets. Box and Whisker plots for select transcripts for control, PAB, and rPAB animals including ANFB, ACTS, PRG4, PDE3A, HA1A, and HA1B (B). TV = tricuspid valve, RV = right ventricle, ATL = anterior tricuspid leaflet, PTL = posterior tricuspid leaflet, STL = septal tricuspid leaflet.

Principal component analysis (PCA) (Figure 5A) demonstrated clear clustering of the control PAB and rPAB RV and TV samples with minimal clustering based on leaflet anatomical locations. Differential gene expression analysis (Figure 5B) revealed 278 differentially expressed genes in the RV between control and PAB, 131 between control and rPAB, and 110 between PAB and rPAB with the TV having 71 between control and PAB, 161 between control and rPAB, and 48 between PAB and rPAB. Log2 FC of control vs PAB and control vs rPAB (Figure 5C) revealed 64 transcripts specifically upregulated in the PAB RV and 17 upregulated in the rPAB RV. One transcript (NAV2) demonstrated upregulation in both the PAB and rPAB RV, while 151 transcripts where downregulated in both PAB and rPAB samples. The TV had 19 transcripts identified to be upregulated in the PAB, 17 in the rPAB, and 117 downregulated in both the PAB and rPAB valves. Gene ontology (GO) enrichment analysis (Figure 5D) of the transcripts upregulated in PAB RV revealed enrichment of fibrinogen and fibrosis, while rPAB analysis showed enrichment in base excision repair. Transcripts downregulated in both PAB and rPAB RV were associated with abnormal cardiovascular systems. GO analysis of the TV did not show any enrichment in the transcripts specific to the PAB and rPAB samples, but the transcripts downregulated in both the PAB and rPAB samples were enriched in liver, bronchial epithelial cell, and occipital lobe.

**Figure 5.**
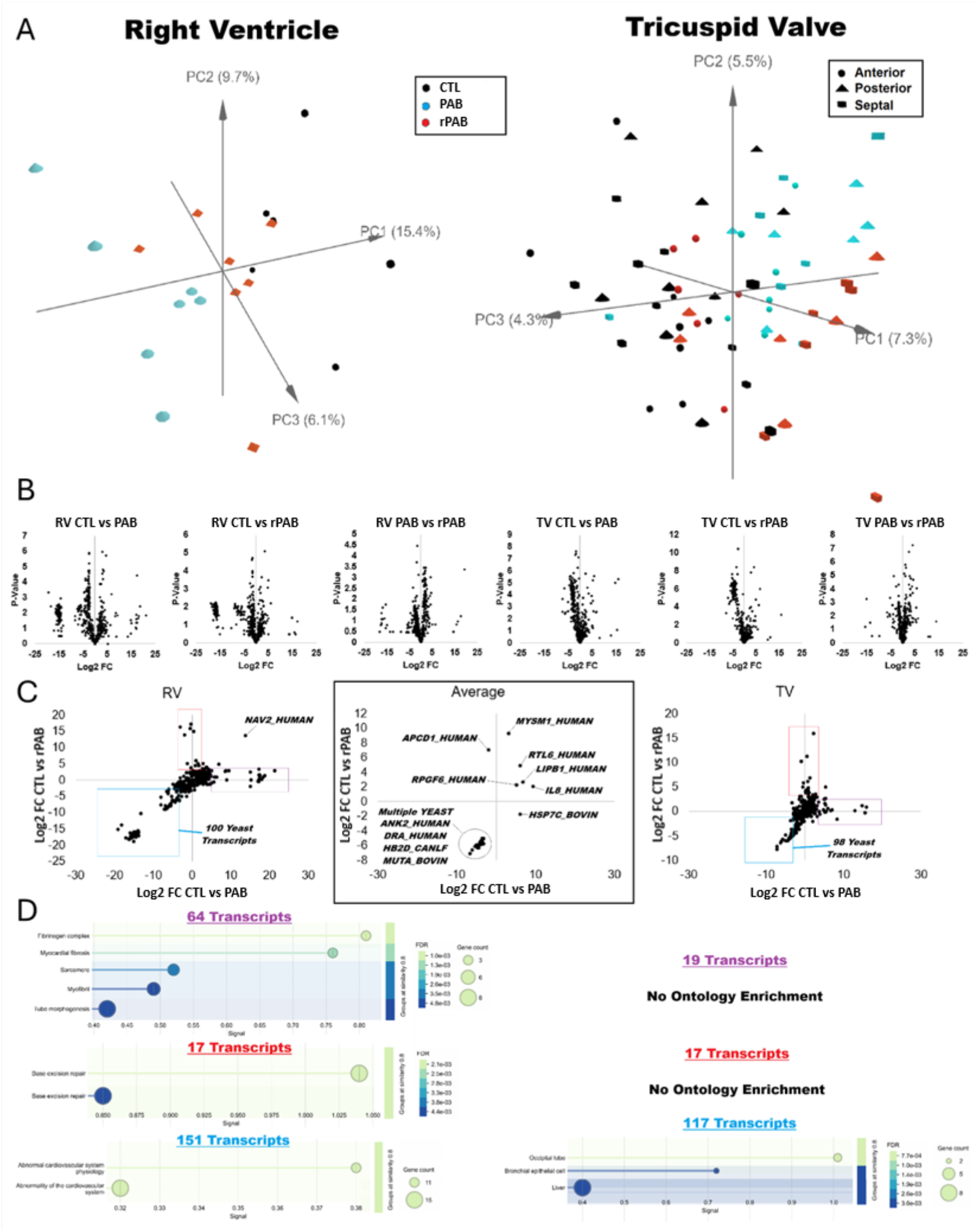
Principal component analysis plots for RNA sequencing data for right ventricle and tricuspid valves (A) demonstrating clustering of samples based on control PAB and reversed status and anatomical location. Volcano plots of differentially expressed genes comparing right ventricle and tricuspid valve tissues between control PAB and rPAB animals (B). Scatter plots identifying high log2 fold change differences in expression between control and PAB (x axis) and control and rPAB (Y axis) for right ventricle and tricuspid valve tissue (C). Gene ontology enrichment of the transcripts with high fold change differences in control vs PAB animals (purple) control vs rPAB (red) as well as both (blue) (D). TV = tricuspid valve, RV = right ventricle, ATL = anterior tricuspid leaflet, PTL = posterior tricuspid leaflet, STL = septal tricuspid leaflet.

## Discussion

This study demonstrates that reversal of pressure overload induced FTR results in substantial improvement in RV geometry and valve competence but does not restore the TV to a healthy state. Although inflammatory and proliferative responses resolved following debanding, leaflet enlargement persisted and was accompanied by distinct extracellular matrix and transcriptomic alterations. These findings indicate that TV reverse remodeling is incomplete and suggest that leaflet biology may contribute to residual or recurrent TR despite correction of the underlying hemodynamic insult.

The extent of RV reverse remodeling after clinical TV repair is strongly associated with favorable long-term survival^16^ and the degree of reverse remodeling, such as improvements in RVFAC and reductions in RV size, helps stratify patients into risk categories.^17^ Efficacy of FTR repair is strongly correlated with RV reverse remodeling, with superior procedural results when residual TR ≤1+ compared to patients with ≥2.^18^ Residual and recurrent TR remain important limitations of contemporary surgical and transcatheter edge-to-edge and replacement therapies.^19,20^ Although recurrence is often attributed to progressive annular dilation and RV remodeling, our findings suggest that leaflet remodeling may represent an additional contributor to recurrent dysfunction. Surgical consideration for FTR focuses on annular dilation and right heart remodeling with leaflets believed to play a passive role; however, recent studies have shown FTR induced molecular and histological changes present within the TV.^12^ This study builds upon this work, demonstrating significant molecular and histological changes in the TV following reverse remodeling.

In clinical practice, patients with FTR secondary to PH experience improvement of regurgitation following treatment, commensurate with resolution of mPAP,^21,22^ leading many to conclude that TR is completely driven by RV geometric changes and annular dilation associated with PH or atrial fibrillation, obscuring the role of and ramifications for the valve itself. This has been supported by the observation of FTR resolution and cardiac remodeling with successful treatment of atrial fibrillation;^23,24^ however, the long-term behavior of the valve tissue in these patients remains poorly understood, with data demonstrating the remodeling of the TV limited to pre-clinical models.^12,15,25–26^ The association of TR with increased rates of cardiovascular-related deaths in patients with heart failure,^27^ and data suggesting reduced short and long-term survival rates after surgical TV intervention,^28^ mandates further investigate the role of the TV leaflets in the pathogenesis of FTR. Our data demonstrates that reverse remodeling did not represent a simple return toward the healthy state. Instead, unloaded leaflets demonstrated structural, histologic, and transcriptional features distinct from both control and pressure- overloaded valves. These findings suggest that TV adaptation establishes a unique biologic state that persists beyond normalization of loading conditions and may influence future valve performance.

Consistent with previous studies leveraging the PAB model,^12,15^ we observed histologically that cross sectional leaflet area significantly increased in PAB animals, and we demonstrated that this increase remained in rPAB animals, indicating that the geometric adaptation of the valve persists even after unloading. While the valves maintained an increased size, the ECM composition of rPAB animals was significantly different than both control and PAB animals, notably in mucin and fibrin content. These findings suggest a shift from matrix expansion to matrix stabilization or repair, potentially mediated by ongoing low-grade inflammation or mechanical stress. These thoughts are corroborated by the finding that immune cells were significantly increased in PAB animals along with proliferative index as measured by Ki67. Interestingly, these changes in immune infiltrate and proliferation were no longer evident following removal of the band, even though the ECM was significantly different than both diseased and healthy states. This indicates that while the active response of the leaflets to pressure overload may be transient, this active maladaptive response leaves behind a wake of a severely altered valve.

Transcriptomic analyses supported the histologic findings by demonstrating persistent alterations in pathways associated with immune regulation, extracellular matrix remodeling, and tissue homeostasis despite normalization of hemodynamic loading. Several transcripts involved in tissue homeostasis and remodeling, including *PRG4* and *PDE3A*, remained dysregulated after debanding, suggesting persistent biological adaptation despite recovery of valve competence and right ventricular remodeling. While *PRG4* has previously been implicated in human valvular disease, particularly calcific valve pathology,^29^ our findings suggest a broader role in valve adaptation to altered mechanical loading. Persistent alterations were also observed in immune pathways, including major histocompatibility transcripts (*HA1A* and *HA1B*) and previously identified inflammatory mediators such as *CXCL8*.^12,30^ Notably, these changes persisted despite normalization of inflammatory cell infiltration and proliferative activity, indicating that recovery of leaflet biology does not parallel recovery of hemodynamics. The persistence of both structural and transcriptional abnormalities following unloading suggests that valve tissue may retain a form of biologic memory of prior mechanical stress. This phenomenon may help explain why restoration of favorable hemodynamics does not always result in complete normalization of valve structure or durable valve competence. Collectively, these data support the concept that FTR is not solely a disease of annular and ventricular geometry but also involves lasting biological changes within the leaflet itself. Such persistent remodeling may represent an underrecognized determinant of residual or recurrent TR following surgical or transcatheter edge-to-edge repair or valve replacement. Consistent with the leaflet findings, gene expression of the RV failed to return to a normal state following debanding. Removal of hemodynamic stress was marked by persistent dysregulation of genes associated with cardiac stress response, remodeling, and tissue repair including *PDE3A*, *NAV2*, *ANFB*, and *ACTS*.^31–34^ This suggests that reverse remodeling of both the ventricle and valve is incomplete despite normalization of hemodynamics. Importantly, however, the response of the RV was distinct from that observed in the leaflet tissue, highlighting differential adaptation of myocardial and valvular structures to pressure overload and subsequent unloading. Together, these data support the concept that cardiovascular tissues retain evidence of prior mechanical stress even after apparent functional recovery.

TV repair strategies primarily address annular geometry, yet durable valve competence depends on the interaction among annular dimensions, leaflet area, tethering forces, and ventricular geometry. Persistent leaflet remodeling may therefore influence repair durability even when annular dimensions and right-heart loading conditions improve. As transcatheter tricuspid interventions continue to expand, including edge-to-edge repair, a more complete understanding of leaflet biology may become increasingly important. Therefore, long-term procedural success is likely determined not only by ventricular and annular remodeling but also by the biological state of the leaflet tissue at the time of intervention.

Adaption of leaflet tissue to hemodynamic stress is a complex process resulting in alterations in many homeostatic processes by the diverse milieu of leaflet cells.^35–37^ When the hemodynamic stress is removed, the tissue again responds resulting in a novel valve phenotype dissimilar to both a homeostatic and a stressed valve. Collectively, these findings challenge the traditional view that FTR is solely a consequence of reversible right-heart geometry. Although relief of pressure overload improved ventricular function and valve competence, leaflet remodeling persisted at structural and molecular levels. These observations suggest that the tricuspid leaflet is an active participant in disease progression and recovery, representing a potentially underrecognized determinant of residual or recurrent TR after surgical or transcatheter intervention. Future studies should determine whether persistent leaflet remodeling influences repair durability and whether valve directed therapies can enhance long-term outcomes.

### Limitations

Though our study was able to generate clinically relevant significant TR, our study is limited to the translation of data generated in animal models not always reflective of human physiology. However, the ovine model represents a clinically relevant model with similar weight and cardiovascular physiology to human subjects. Our study attempted to model the resolution of PH through removal of the PAB inducing FTR, and though pressure overload was resolved, volume does not simultaneously return to baseline, limiting conclusions on causality in the relationship between pressure and volume overload and reverse remodeling of the TV. In addition, our study was limited to a single timepoints (8 weeks) of PAB induced FTR and PH, as well as a single timepoint of reverse remodeling following removal of the PAB (8 week after removal of the PAB). Due to this limitation, we are unable to resolve if histologic and molecular alterations have reached full resolution or if our observations are part of the trajectory of resolution.

## Nonstandard Abbreviations and Acronyms

FTR: functional tricuspid regurgitation
LV: left ventricle
LVEF: left ventricular ejection fraction
LVEDV: left ventricular end diastolic volume
ECM: extra cellular matrix
PA: pulmonary artery
PAB: pulmonary artery banding (banded group)
PRG4: proteoglycan 4
rPAB: reversed pulmonary artery banding (debanded group)
PH: pulmonary hypertension
RA: right atrium RV right ventricle
RVEDV: right ventricular end diastolic volume
RVFAC: right ventricular fractional area change
sPAP: systolic pulmonary artery pressure
TA: tricuspid annulus
TAPSE: tricuspid annular plane systolic excursion
TR: tricuspid regurgitation
TV: tricuspid valve

---

**Supplemental Table 1.** Echocardiographic parameters at 8 weeks in interventional groups.

| Parameters at 8 weeks | PAB | rPAB | p-value |
| --- | --- | --- | --- |
| HR (beats/min) | 120 ± 35 | 135 ± 19 | 0.286 |
| Weight (kg) | 79 ± 3 | 76 ± 5 | 0.171 |
| TR (0-4+) | 3.7 ± 0.5 | 3.2 ± 1 | 0.178 |
| TA (cm) | 3.9 ± 0.4 | 4 ± 0.6 | 0.530 |
| RA area (cm) | 29 ± 5 | 23 ± 5 | 0.037* |
| RVFAC (%) | 29 ± 13 | 21 ± 8 | 0.212 |
| LVEF (%) | 53 ± 4 | 60 ± 7 | 0.250 |
| MR (0-4+) | 0.5 ± 0.5 | 0.5 ± 0.7 | 1.00 |

